# Neuroimage Denoiser - a Deep Learning Framework for Removing Noise from Transient Fluorescent Signals in Functional Imaging

**DOI:** 10.1101/2024.06.08.598061

**Authors:** Stephan Weißbach, Michela Borghi, Jonas Milkovits, Carolina Amaral, Abderazzaq El Khallouqi, Susanne Gerber, Martin Heine

**Affiliations:** Institute of Developmental Biology and Neurobiology (iDN), Johannes Gutenberg-University, 55128 Mainz, Germany; Institute of Human Genetics, University Medical Center, Johannes Gutenberg-University, 55131 Mainz, Germany

**Keywords:** glutamate imaging, calcium imaging, functional imaging, iGluSnFR, GCaMP6f, denoising, deep learning

## Abstract

We developed Neuroimage Denoiser, a novel U-Net-based model that effectively removes noise from microscopic recordings of transient local fluorescent signals. The model makes the denoising process independent of the recording frequency and the kinetics of the sensor used. The framework is easy to use for denoising and training and has minimal hardware requirements, thus, making it accessible for an average laboratory to create a custom version specific to their experimental setup. Neuroimage Denoiser significantly enhances the quality of functional microscopy recordings by effectively removing noise, thereby facilitating a more accurate and reliable analysis of neural activity.

**Highlights:**

- Neuroimage Denoiser is a deep learning framework to remove noise from functional microscopic recordings, particularly trained and tested for glutamate imaging
- Neuroimage Denoiser balances the removal of noise while preserving the amplitude of responses
- Neuroimage Denoiser operates without re-training for different sensors (when the localization is similar) and recording frequencies

**Motivation:** Accurate measurements of neuronal activity through functional imaging are critical in understanding mechanisms of synaptic plasticity and learning concerning changes in the molecular composition of single synapses. Traditional denoising methods, such as Gaussian or Median filters, indiscriminately smooth entire recordings, reducing temporal and spatial resolutions considerably. Existing frameworks are not suited to remove noise from glutamate recordings due to the fast dynamics of the sensor. Therefore, a specialized tool for the challenges imposed by glutamate recordings, i.e. faster dynamics, and synaptic localization, is needed.

## Introduction

Noise is a pervasive phenomenon impacting all experimental systems used in neuroscience. Accurate measurement of neuronal activity is crucial for understanding neuronal function, yet noise in imaging systems can severely distort data^1^. With the development of genetically encoded calcium or neurotransmitter sensors, the possibility reading the activity of neuronal subcellular compartments like synapses became feasible, based on fast time-lapse fluorescent imaging. In camera-based setups, noise affects pixels individually without any spatiotemporal relations, often uneven distributed over the field of view. However, the signal from fluorescent reporters that indicates neuronal activity in synapses or somata exhibits a high spatiotemporal correlation.

Traditional methods, such as Gaussian or Median filters, can be used to remove noise from microscopic recordings. While effective to some extent, these filters indiscriminately smooth the entire image, leading to reduced temporal and spatial resolutions^2^. Additionally, the manual selection of filter parameters can be labor-intensive and subjective, often resulting in suboptimal noise reduction.

There has been a considerable effort to remove noise from calcium imaging data and several frameworks have been proposed. Current denoising approaches for fluorescence imaging largely rely on temporal correlations in the signal and are optimized for large somatic structures. DeepInterpolation^3^, developed for somatic GCaMP6f imaging, uses frame interpolation between time points (similar to Noise2Void^4^ and Noise2Self^5^). DeepCAD^6^ and DeepCAD-RT^7^ use a 3D U-Net that is trained on two split sub-stacks consisting of interlaced frames from the raw somatic GCaMP6f recording leveraging the spatio-temporal features of signal in comparison to noise. Training 3D U-Net can be challenging since the high amount of parameters allows overfitting easily^6^. DeepSeMi^8^, though more versatile with multiple organelle markers, also incorporates temporal information through its hybrid spatial-temporal network. SUPPORT^9^ integrates spatial and temporal information to denoise a range of fast voltage sensors that show a somatic localization. However, the authors show its applicability to other functional imaging sensors such as GCaMP8f and also conduct limited tests for the synaptic iGluSnFR and iGABASnFR. Recently, Wang and colleagues introduced CellMincer^10^ a two-stage denoising framework consisting of a frame-wise 2D U-Net combined with a pixel-wise 1D convolutional model to capture spatial and temporal dimensions respectively. CellMincer is specifically trained to denoise fast voltage sensor recordings.

Previous approaches mostly share two key limitations. First, they are primarily optimized for large cellular structures like somata rather than synapses. Second, their dependence on the temporal dimension necessitates retraining for different recording frequencies and sensor combinations, which demands substantial data collection and computational resources for each new experimental condition.

Commonly used sensors like somatic GCaMP6f, which measure calcium influx, exhibit slower dynamics compared to diffusion-dominated local release of neurotransmitters. Thus, monitoring synaptic glutamate release requires considerably higher spatial and temporal resolution^11^—the glutamate sensor iGluSnFR3.v857.SGZ exhibits orders of magnitude faster kinetics than somatic calcium indicators used in most previous studies, with signals originating from tiny synaptic release sites orders of magnitude smaller than somata. This combination of rapid dynamics and synaptic localization presents unique challenges: the signals are spatially confined, temporally sparse, and accompanied by higher background noise compared to somatic recordings.

While existing denoising methods excel at enhancing slow somatic signals by leveraging their temporal continuity, they may introduce artifacts when applied to rapid synaptic events that appear discontinuous between frames. The need to retrain models for each combination of sensor and recording frequency further complicates their application to diverse imaging conditions. Therefore, a more flexible approach that can adapt to different temporal dynamics without retraining and specifically addresses the spatial scale of synaptic signals is needed to preserve the fast dynamics and spatial precision of synaptic glutamate imaging.

To address these specific challenges, we developed Neuroimage Denoiser, a novel model that effectively removes independent noise from microscopic recordings. Unlike many recent methods, the training and architecture of Neuroimage Denoiser makes the denoising process independent of the recording frequency and the kinetics of the sensor used. This flexibility sets Neuroimage Denoiser apart from previous methods and makes it a versatile tool for various experimental setups, including fast calcium imaging.

Neuroimage Denoiser has minimal hardware requirements, making it accessible for many laboratories to create a custom version tailored to their specific experimental setup. Neuroimage Denoiser enables researchers to obtain high-quality, noise-reduced data without the need for extensive computational resources or expertise.

## Material and Methods

### Primary hippocampal mouse cultures

Co-authors of the paper acquired the data used for different projects. The use of mice in research was approved by District administration Mainz-Bingen, 41a/177-5865-§11 ZVTE, 30.04.2014. Primary mouse hippocampal cultures were generated from newborn pups^12^. Animals were sacrificed according to the European and local government (Rheinland-Pfalz, Germany) regulations for animal welfare. Briefly, newborn pups of either sex were anesthetized on ice and sacrificed using the decapitation method. Following hippocampal isolation, cells were dissociated with trypsin (Thermo Fisher Scientific; Cat#: 15090-046) and plated on poly-L-lysine-coated 18 mm glass coverslips with a density of 70,000 cells per coverslip. After 1 h incubation in Dulbecco’s Modified Eagle Medium (Thermo Fisher, Gibco; Cat#: 41966-029/D6429) supplemented with 10% fetal bovine serum and 0.5% glutamate at 37°C, coverslips were transferred to 12-well plates containing 1mL of growth medium composed of Neurobasal-A (Thermo Fisher, Gibco; Cat#: 12349-015) with 2% GlutaMax (Thermo Fisher, Gibco; Cat#: 35050-038), 2% B27 (Thermo Fisher, Gibco; Cat#: 17504-044), 0.1 M sodium pyruvate (MERK, Sigma; Cat#: S8636). Cultures were maintained in a humidified incubator with an atmosphere of 95 % air and 5 % CO_2_ at 37°C.

### Neurons transfection

To perform functional imaging experiments, neurons were transfected at DIV3-5 using the calcium phosphate method with either the glutamate sensor iGluSnFR3.v857.SGZ (Addgene; Cat#: 178330)^13^, iGluSnFR.S72A (Addgene; Cat#: 106176)^14^, or the calcium sensor pAAV-hSyn-synaptophysin-GCamp6f (cloned by Dr. Arun Chhikara from Addgene plasmid Cat#: 67634 and phSyn-Synapthophysin-GCamp6f). Briefly, the neuron’s growth medium was exchanged with 1 mL of pre-warmed Neurobasal-A medium, and the original medium was stored and kept at 37°C. In the meantime, per 18 mm coverslip, 30 µL of transfection buffer containing in mM: 274 NaCl, 9.5 KCl, 1.4 Na_2_HPO_4_, 15 Glucose, 15 HEPES, pH 7.14 were added to a solution containing 3 μg of DNA, 3 µl of CaCl_2_ (2M) and 24 µL of H_2_O (q.s. 30 µL). The resulting DNA/CaPO_4_ mix was incubated for 20 min at RT and then added in a proportion of 60 µl of mix per well. Incubation on neurons was maintained at 37°C until precipitate formation (40 min). Following DNA/CaPO_4_ incubation, neurons were washed two times for 5 min with 1X HBS buffer containing in mM: 137 NaCl, 4.75 KCl, 0.7 Na2HPO4, 7.5 Glucose, 7.5 HEPES, pH 6.9 and for one time for 5 min with Neurobasal-A medium. Following the washing steps, the original growth medium was added, and neurons were stored in the incubator at 37°C until live imaging experiments were performed at DIV16 - 19.

### Functional Imaging

All live experiments were conducted on hippocampal cultures at DIV16 - 19 at 37°C within a whole microscope incubator. Neurons were perfused with extracellular solution containing (in mM): 145 NaCl, 2.5 KCl, 10 HEPES, and 10 D-glucose 2 MgCl_2_, 2 CaCl_2_ (pH 7.4). Network activity was suppressed by adding 10 μM CNQX (Tocris; Cat#: 1045) and 10 μM D-AP5 (Tocris; Cat#: 0106) to the extracellular solution as indicated for the respective experiment. Experiments were performed with an inverted total internal reflection fluorescence (TIRF) setup composed of the eclipse Ti microscope (Nikon) equipped with a 60 x Apo TIRF oil objective (1.49 NA; Nikon). Fluorescence was excited with a combined laser system (Coherent; MPB Communications Inc.). Images were captured by a scientific CMOS camera (ORCA-Fusion C14440-20UP, Hamamatsu) at frame rates between 100 to 1000 Hz controlled by NIS-Elements Advanced Research acquisition software (Nikon). Electrical field stimulation was carried out using two platinum electrodes placed on the coverslip at a 10 mm distance. For glutamate imaging experiments, triggered electrical field stimulation was used to elicit 20 action-potential-like stimuli. The extracellular stimulus sequences (0.9 ms; 50 mA; 1 s interval) were triggered by an isolated pulse generator (A&M Systems, Model 2100) and delivered via a stimulus isolator unit A385 (WPI). Images were acquired by image streaming for 2000 frames within a defined region of interest (ROI) 800 × 800 pixels (100 Hz acquisition rate) or 250 × 250 pixels (1000 Hz acquisition rate) with a pixel size of 72nm. For calcium imaging experiments, each region of interest of 512 × 512 pixels was stimulated using 1,3 and 5 action potential-like stimuli (1 ms, 50 mA, 20 ms interval).

### Synaptic ROI and Threshold

Circular synaptic ROIs were selected manually using Fiji (version 1.54f)^15^ and set to a diameter of 9 pixels. The average traces along the z-axis of the synapses were extracted by either Fiji (version 1.54f)^15^ or Python (version 3.10.14)^16^ with roifile (version v2024.3.29)^17^.

On the average traces, the threshold was computed using numpy (version 1.26.4)^18^ based on the baseline frames up to the first stimulation, using the formula:

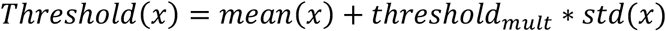

Whenever the average trace crossed the computed threshold (set by threshold_mult_), this was considered to be a response.

### Signal-to-noise ratio

To assess the denoising performance, we calculated the signal-to-noise ratio (SNR) for both raw and denoised glutamate and calcium recordings. SNR was computed for each synaptic response using the mean traces from synaptic ROIs according to the formula:

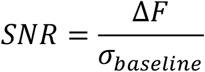

where ΔF represents the signal amplitude (calculated as the difference between the peak intensity value and mean baseline), and σ_baseline_ is the standard deviation of the baseline. The baseline was defined as the ten frames preceding the initial threshold crossing. Statistical comparison between raw and denoised recordings was performed using a two-tailed paired t-test using scipy (version 1.14.1)^19^.

We further compared the single-pixel SNR between the raw and denoised recording, which was defined as the mean fluorescence value divided by the standard deviation along the temporal axis using numpy (version 1.26.4)^18^. Subsequently, we randomly sampled 10,000 pixels and plotted the distribution with a seaborn histplot (version 0.13.2)^20^.

### Preparing Training Data

Training deep learning models is sensitive to imbalances between foreground and background^21,22^. Different from synaptic GCaMP6f recordings used for the training of several previously described denoising methods^3,6,7^, the microscopic recording of synaptic glutamate responses is dominated by background due to the small synapses that sparsely respond when defining a responding synapse as foreground. Therefore, the available data is filtered for active regions and frames to remove the background (Figure 1a). We transformed our recording frames into activity maps to filter efficiently. First, the data is transformed by a pixel-wise rolling window (along 50 frames) z-normalization (Figure 1b). The rolling window adapts for potential bleaching in the recording.

**Figure 1.**
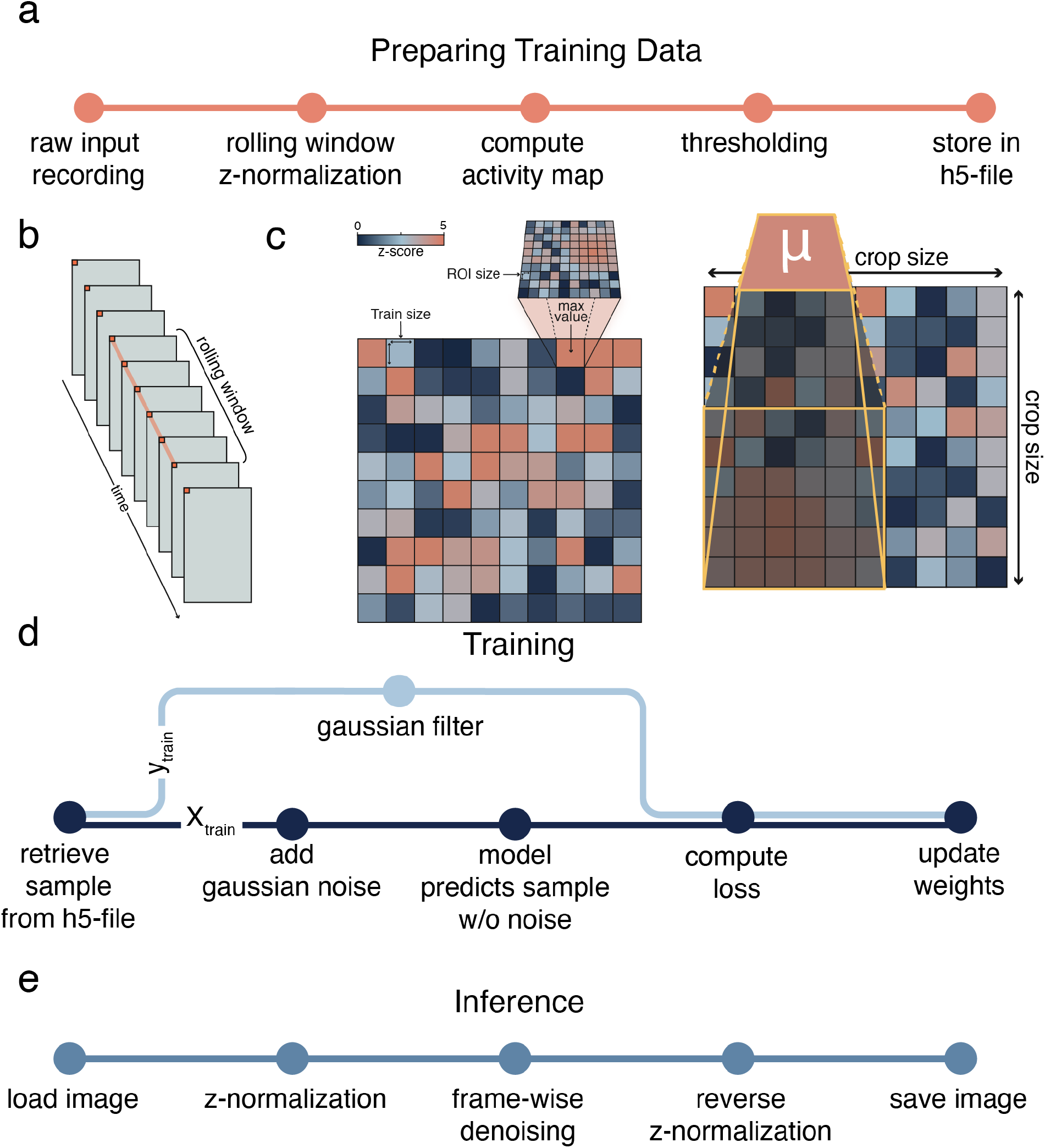
a) Schematic representation of the preparation of the training data. The preparation aims to enrich the foreground (crops of the recording with responding synapses/somata) over the background (non-responding frames or crops) and store them in an h5 file for training. b) Z-normalization for training data preparation is computed per pixel using a rolling window to account for potential bleach occurring during the recording. c) Schematic representation of an activity map that is computed for each frame of a recording during the training data preparation step. The frame is divided into crops of equal size. Per crop, the average z-score per possible ROI of a given size is computed and whenever this averaged z-score exceeds a set threshold, the crop is added to the training data. Left: one frame divided into crops and applied average filter; right: average filter applied of ROI size 6 to a crop d) Schematic representation of the training routine. X_train_ represents the input on which the model makes its prediction, and the result is compared to y_train_. f) Schematic representation of the inference/denoising process. The inference is done on a z-normalized representation of the recording, which is reversed after denoising.

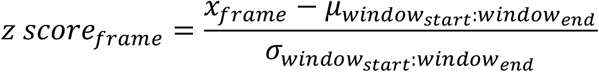

An activity map is calculated framewise and represents the maximum z-score within a local region for each spatial position in each frame. Each frame is divided into crops of a predefined size of 32 × 32 pixels to compute the activity map. A crop is assigned with the maximum value of the kernel-averaged region, obtained by sliding a quadratic kernel over the crop (step size 1) and computing the average intensity within that kernel at each position (Figure 1c, right). The kernel size corresponds to the expected minimal region of interest (ROI) size, reflecting the spatial extent of the structures under investigation (6 pixels for synapses in our setup). The activity map effectively captures and highlights regions exhibiting high-intensity fluctuations that are potential indicators of neuronal activity, by assigning the maximum averaged kernel value to each crop. Next, these activity maps are filtered for crops that exceed a minimum z-score of 3. Subsequently, the pixel-wise z-score (without using a rolling window) is computed, and the transformed crop is stored in an h5 file for training, using h5py (version 3.11.0).

Neuroimage Denoiser can be run on an entire directory storing multiple recordings and will automatically prepare the training data without manual intervention. Additionally, a memory-optimized version is implemented that necessitates less RAM at the cost of a longer run-time.

### Model Architecture and Training

The U-Net model^23^ is implemented in PyTorch (version 2.1.1.)^24^ with a single input and output channel (Figure 1a). The U-Net architecture consists of an encoder and a decoder. The encoder comprises blocks containing two 2D convolutional layers with 3×3 kernels, ReLU activation, and batch normalization, followed by a 2D max-pooling layer for downsampling. These blocks are referred to as down-blocks. Skip connections connect the corresponding down-blocks to the up-blocks in the decoder without any processing.

The decoder comprises up-blocks, each consisting of a 2D transpose convolutional layer for upsampling, followed by a concatenation with the respective feature map from the corresponding down-block (skip connection). This concatenated output is then processed by two 2D convolutional layers with 3×3 kernels, ReLU activation, and batch normalization. The final layer is a 1×1 2D convolutional layer that maps the feature vector to the denoised image. This architecture allows the network to capture both local and global features, enabling accurate denoising of the input image. Using a fully convolutional model without dense layers allows flexible input image sizes during inference, limited only by GPU memory, overcoming fixed training input constraints.

Training is performed on z-transformed patches containing active synapses for one epoch using a batch size of 64, the ADAM optimizer with a learning rate of 0.0001, on a consumer-grade NVIDIA GeForce 2060 super graphics card with 8GB of vRAM. A grid search for the parameters listed in Table 1 was performed to find a suitable combination of parameters for the training. The model’s objective is to reduce noise in the recordings while maintaining the integrity of the peak response amplitudes.

**Table 1.**
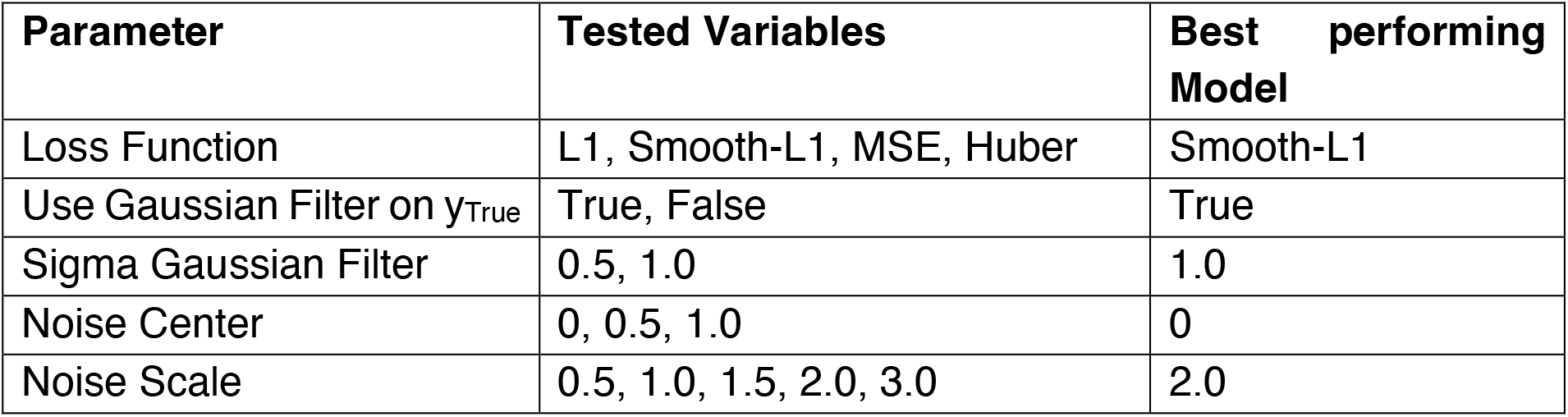
Model Optimization Parameters: This table presents the various parameters tested during the optimization of the Neuroimage Denoiser model. For each parameter, different variables were tested to determine the most effective configuration. The “Best performing Model” column lists the values that yielded the best performance based on the evaluation metrics.

To evaluate the model’s performance in preserving the amplitude of neural responses, regions of interest (ROIs) exhibiting clear stimulus-evoked responses were manually identified in the raw recorded data. The peak response amplitudes within these ROIs were measured following the presentation of a stimulus. After training the model, the same ROIs were analyzed in the denoised output, and the corresponding peak response amplitudes were extracted. The model’s accuracy in preserving the response amplitudes is quantified by calculating the Mean Absolute Percentage Error (MAPE) using scikit-learn (version 1.5.0)^25^ between the raw and denoised peak amplitudes across all ROIs. The correlation between raw and denoised peak amplitudes was calculated using scipy (version 1.14.1)^19^.

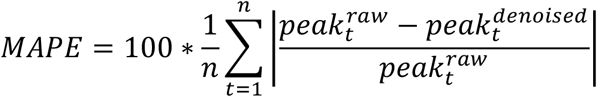

Additionally, it is crucial that the model efficiently removes noise from the microscopic recordings. This is measured by the standard deviation of non-responding sections of the recordings.

Both metrics (MAPE and standard deviation) are ranked, and a combined score is calculated. The best-performing model is selected based on the minimal combined score achieved (supplementary table 1).

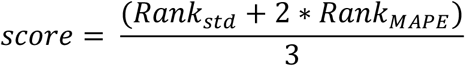

### Inference/Denoising

The denoising (or inference) using Neuroimage Denoiser can be done on single images or entire directories containing multiple sub-directories while maintaining the original folder structure. The process is schematically visualized in Figure 1e. First, the input image is loaded in tiff or nd2 format using tifffile (version 2023.12.9)^26^ or nd2 (version 0.8.1) respectively and a z-normalization (along the time axis; no rolling window) is applied. The mean and standard deviation per pixel are temporarily saved. Neuroimage Denoiser performs the denoising on the z-normalized frames and afterward, the resulting frames are transformed back into intensity values using the saved mean and standard deviation. The denoised image is saved as a tiff file using tifffile (version 2023.12.9)^26^.

### Reimplementation of DeepInterpolation

To fairly compare the performance of DeepInterpolation^3^ with the approach of Neuroimage Denoiser, we re-implemented the model in PyTorch (version 2.1.1)^24^. Therefore, the model can be trained on crops of active synapses (32 × 32 pixels) and comparability was ensured. The number of frames before and after the target frames was reduced to five frames to account for the faster dynamics of glutamate sensors (supplementary Figure 1a). The model was trained using the ADAM optimizer with an L1 loss function, and a learning rate of 0.0001 on an NVIDIA GeForce GTX2060 super graphics card.

### Data Visualization

Data was visualized using Python (version 3.10.14)^16^, matplotlib (version 3.9.0)^27^, and seaborn (version 0.13.2)^20^. Schematic drawings were made using Adobe Illustrator.

## Results

Denoising glutamate imaging data presents unique challenges compared to calcium imaging, as glutamate sensors are orders of magnitude faster and the synaptic localization results in considerably less foreground signal relative to somatic calcium recordings. Here, we present Neuroimage Denoiser, a U-Net-based imaging denoising tool (Figure 1a) specifically trained to remove noise from microscopic glutamate time-series recordings.

To curate a balanced training dataset adjusted to the challenges posed by glutamate imaging data, the recordings are automatically screened to identify frames and regions exhibiting active synaptic signals (Figure 1a) in the training preparation step without manual intervention. This screening can be adapted to different ROI sizes representing the size in pixels of the biological structure where the sensor is localized. We set the ROI size to 6 pixels to account for the size of synapses harboring the sensor in our microscopic set-up.

Initially, we explored training a model in the style of DeepInterpolation^3^. DeepInterpolation predicts a target frame based on a set number of frames preceding and succeeding the target frame (n_pre_, n_post_)^3^. To account for the fast dynamics of the iGluSnFR3 sensor compared to Gcamp6f, we set n_pre_ = 5 and n_post_ = 5 and the model predicts the target frame in the middle (supplementary Figure 1a). However, this temporal interpolation approach, while effective for slower calcium dynamics, did not adequately preserve the fast glutamate response and led to artifacts in the amplitudes (supplementary Figure 1b).

Given the limitations of an approach that leverages the temporal dimension for denoising for capturing these rapid synaptic signals, we turned to the Noise2Noise concept proposed by Lehtinen and colleagues^28^. Their work demonstrated image restoration without ground truth data by training networks to distinguish signals from noise patterns by adding artificial noise to noisy images. Further, this simple approach was the best-performing approach in the Fluorescence Microscopy Denoising dataset benchmarking, outperforming other methods including DnCNN by almost 2dB in PSNR^29^. Leveraging this approach, Neuroimage Denoiser is trained according to the Noise2Noise^28^ method on individual frames and subsequently circumvents the dependence on the recording frequency or the dynamics of the sensor. Therefore, a trained model can be applied to a broad range of experimental set-ups independently of the recording frequency and sensor dynamics.

We conducted a grid search to identify the optimal combination of training parameters (Table 1) and selected the model that best balanced noise removal and amplitude preservation (supplementary Table 1). The best-performing model was trained using a smooth L1-loss. The training data included Gaussian noise added with a noise center of 0.0 and a noise scale of 2.0. Notably, higher noise scale and center values yielded models that removed noise more effectively but at the cost of less reliable amplitude preservation (supplementary Table 1). The selected model effectively removes noise from the raw recording, both visually as measured in the z-profile of the synaptic ROI (Figure 2b), while preserving the amplitudes well (Figure 2c), as evidenced by a significant correlation between raw and denoised amplitudes (R=0.9966, p < 0.0001). The model’s denoising performance is further characterized by the distribution of single-pixel SNR values (Figure 2d). While raw recordings show a narrow distribution concentrated at low SNR values, denoised recordings exhibit a broader distribution shifted towards higher SNR values, indicating comprehensive noise reduction across the entire field of view. This global improvement in signal quality is also reflected in the significant 5.65 ± 0.33 improved SNR for the peaks (mean ± SEM, p < 0.0001, paired one-sided t-test).

**Figure 2.**
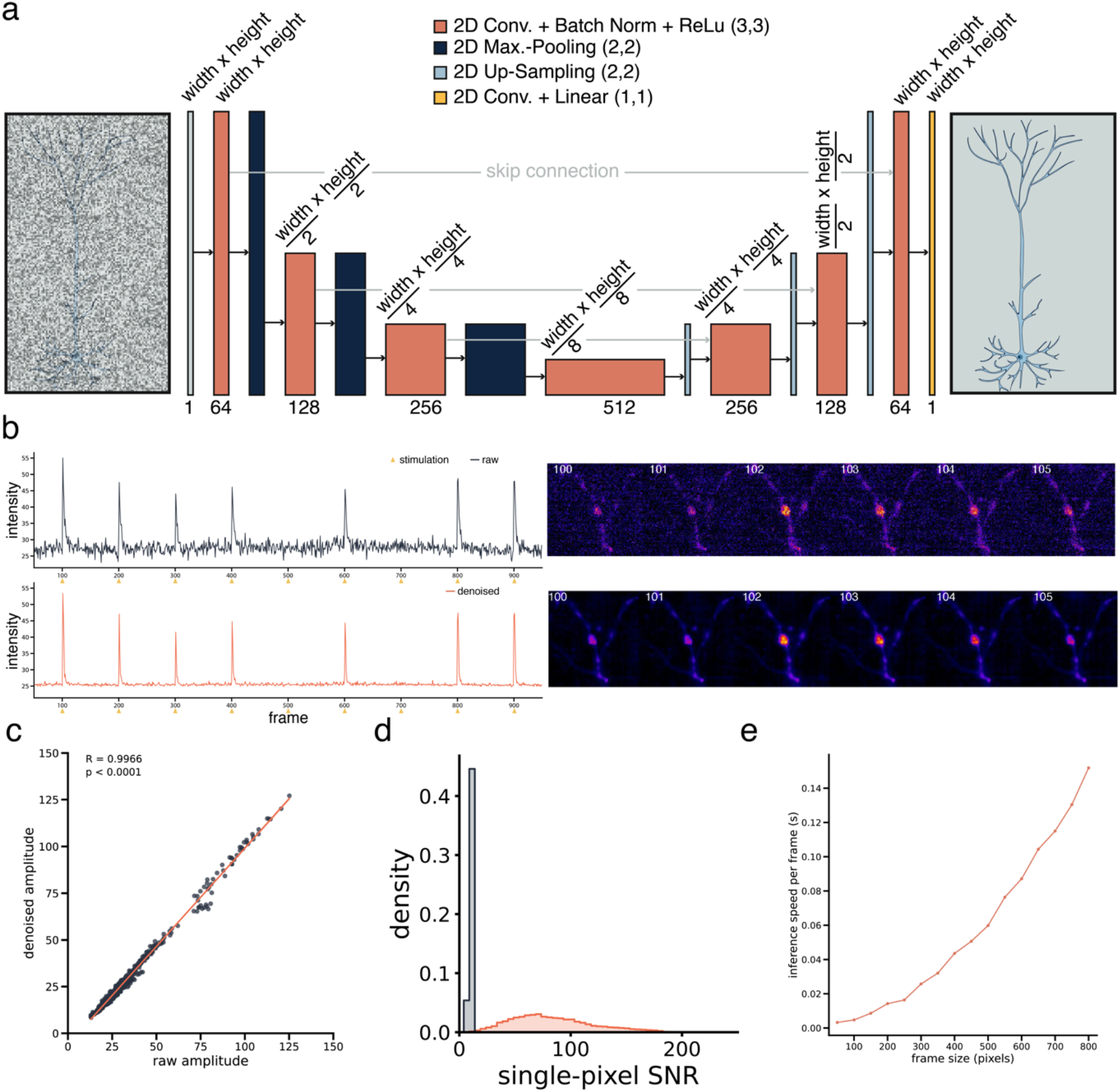
a) Schematic representation of the architecture used for Neuroimage Denoiser is a U-Net^23^ with one input and output channel. The data runs through the encoder and decoder part of the network while skip connections preserve untransformed data. Convolutional layers (orange) use a 3 by 3 filter with ReLu activation. The output layer is a linear layer that does not use an activation function. b) Representative area of a glutamate recording acquired at 100Hz. The stimulus is given in frame 100 and the peak response is in frame 102. After denoising, the amplitude height and time points are well preserved, and the noise level is minimal. c) Neuroimage Denoiser denoised amplitude heights of manually selected ROIs tested against the raw amplitude heights, showing a high correlation (R=0.9966, p<0.0001). d) Single-pixel SNR distributions for raw and denoised recordings. e) Denoising inference speed (frames per second) tested for frame sizes 50 - 800 pixels (averaged values of three repetitions of recordings with 1,000 frames). The image was processed according to Figure 1e)

Neuroimage Denoiser demonstrates fast inference, as tested on three recordings of 1000 frames with varying spatial dimensions (50 to 800 pixels for x and y). The entire process, including normalization and reverse normalization, as shown in Figure 1e, requires between 0.0003 and 0.1519 seconds per frame (Figure 2e). Therefore, denoising 800 × 800 pixels recording of the length of 1,000 frames takes ∼152 seconds.

### Neuroimage Denoiser enables precise Signal Localization across diverse Recording Conditions

Only infrequent spontaneous activity can be observed when recording a neuronal culture in the absence of electric stimulation. Specifically, it is challenging to differentiate the components at complex synapses, which comprise an accumulation of pre-synaptic inputs and post-synaptic receivers. These synapses often involve multiple release sites in close proximity, and the released glutamate can diffuse from its initial release site, leading to highly correlated signals at neighboring post-synaptic sites (Figure 3a, left). However, the maximum localization of a spontaneous release event indicates the release site. Due to the spatial proximity within these complex synapses, identifying and differentiating post-synaptic receiving sites is difficult when the signal is superimposed by noise (Figure 3a, top). After removing the noise using Neuroimage Denoiser, each of the three distinct sites within the complex synapse becomes visible and can be accurately localized (Figure 3a, bottom).

**Figure 3.**
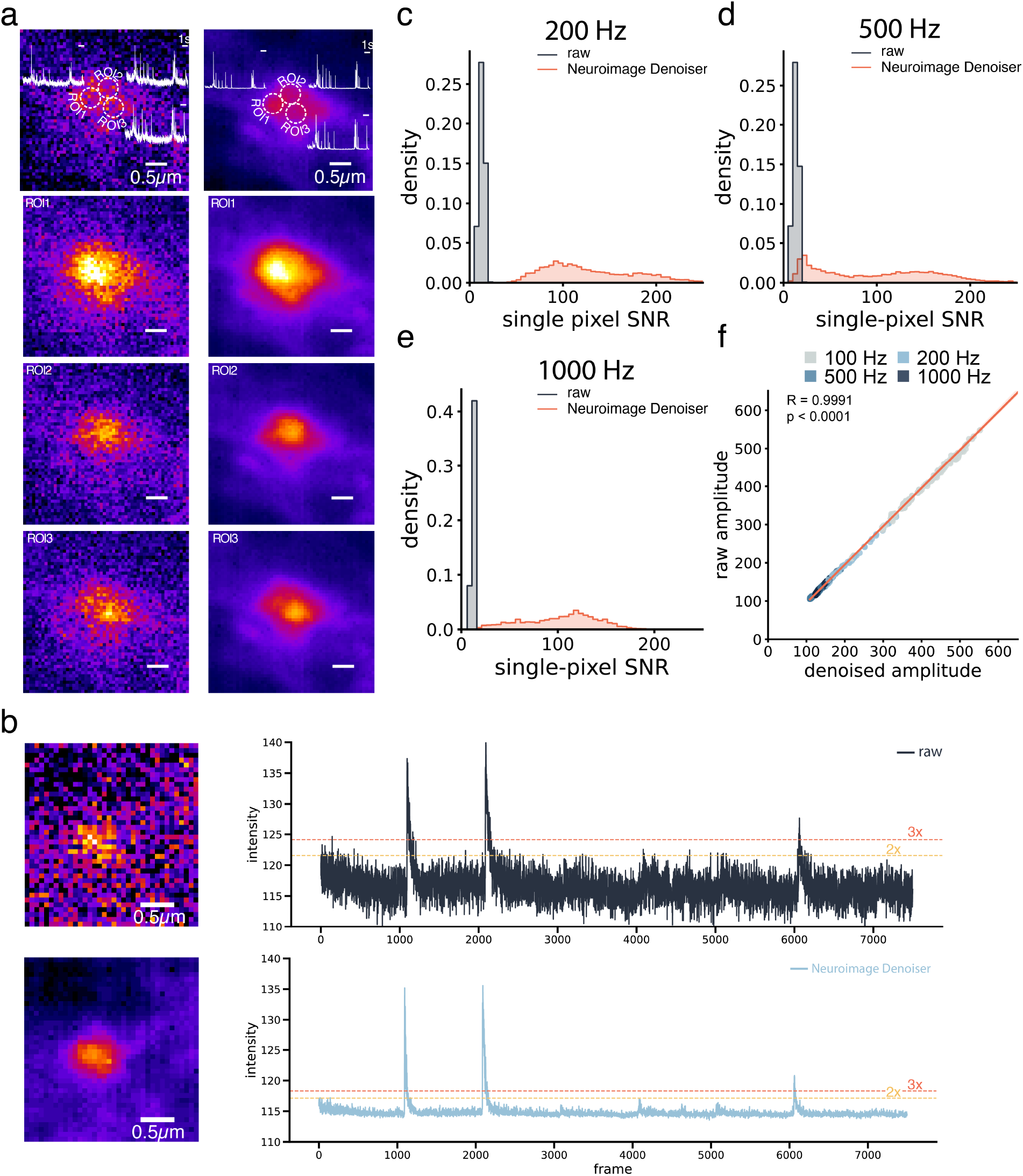
a) Complex synapse with three sites that receive spontaneous input from different pre-synaptic sites. The left column shows the raw recording as it was acquired at the microscope, and the right column shows the same frames denoised by Neuroimage Denoiser. The three different sites showing spontaneous activity are visible. The traces at these sites are highly correlated due to their proximity. b) Neuroimage Denoiser works on recordings acquired with very high recording frequencies (tested here with 1000 Hz), although being trained on 100 Hz recordings exclusively. c-e) Single-pixel SNR distributions for raw and denoised recordings at different acquisition frequencies (200 Hz, 500 Hz, and 1000 Hz). f) Correlation between raw and denoised signal amplitudes, color indicates recording frequency (gray: 100 Hz, light blue: 200 Hz, blue: 500 Hz, dark blue: 1000 Hz)

While Neuroimage Denoiser was trained exclusively on recordings done with 100 Hz, the model is capable of denoising recordings acquired at significantly higher acquisition rates, such as 1000 Hz (Figure 3b). The ability to acquire data at such high frequencies is valuable for capturing rapid synaptic events and amplitudes with high temporal precision. However, high noise levels often complicate these high-frequency recordings, making it challenging to identify true responses amidst false-positive threshold crossings (Figure 3b, top). The denoised recording, on the other hand, has a considerably improved signal-to-noise ratio (Figure 3b, bottom), like the improvement observed in previous recordings at 100 Hz.

Quantitative analysis of single-pixel SNR distributions across different acquisition frequencies (200 Hz, 500 Hz, and 1000 Hz) demonstrates consistent noise reduction capabilities of Neuroimage Denoiser (Figure 3c-e). Further, we also specifically analyzed the SNR for responses at synapses to evaluate the biologically relevant signal improvement. At these functional sites, the denoising significantly improved the SNR with an average 5.21 ± 0.19-fold (mean ± SEM, paired one-sided t-test, p < 0.0001), 7.72 ± 1.18-fold (mean ± SEM, paired one-sided t-test, p < 0.0001), and 8.97 ± 1.18-fold (mean ± SEM, paired one-sided t-test, p < 0.0001) enhancement for 200 Hz, 500 Hz, and 1000 Hz recordings respectively.

Therefore, Neuroimage Denoiser can facilitate the analysis of recordings acquired at very high frequencies without the need to re-train a new model. Acquiring at these high rates opens the door to various applications, such as analyzing multi-vesicular or asynchronous releases from a single synapse, which would yield variable amplitudes and subsequent peaks, or resolving the precise time from stimulus to glutamate release with higher temporal resolution. Additionally, it enables the study of rapid synaptic plasticity mechanisms and the precise kinetics of neurotransmitter release and clearance. The camera-based imaging will allow the comparison of direct multiple synapses in the field of view, which will outcompete scanning microscopy-based approaches, where only single synapses can be imaged at necessary high acquisition rates (>100 Hz).

### Application of Neuroimage Denoiser on alternative Glutamate and Calcium Sensor to study Synaptic Plasticity

Since Neuroimage Denoiser is not bound to a certain sensor and its associated kinetics but only to its localization, we applied the denoising to recordings of fast iGluSnFR.S72A^14^. This version of the sensor has faster kinetics, but a lower signal-to-noise ratio compared to iGluSnFR3.v857.SGZ^13^ particularly due to the high noise level present in the baseline of the recording (until stimulation at frame 100). This noise in the baseline elevates the threshold above which a signal is considered a release event. iGluSnFR.S72A^14^ is useful for plasticity studies since the kinetic properties allow the recording of synaptic release events up to 20 Hz pulse interval. Conventionally such recordings are done via patch-clamp^14^, which will not allow reaching the single synapse resolution as easily as imaging approaches. Conversely, iGluSnFRs timing and localization of excitatory synaptic inputs^11^ enable a more complete picture of the synaptic plasticity of a neuron under a given condition, which is of particular interest for investigating the molecular mechanisms underlying synaptic plasticity. Key plasticity metrics are the paired-pulse ratio, quantifying facilitation or depression of consecutive responses, and the failure rate, measuring the proportion of stimuli failing to evoke a response hence indicating release probability. Nevertheless, due to the low signal-to-noise level and the fast bleaching of the sensor, the recordings are superimposed by noise (Figure 4a, left) that considerably complicates the identification of responding synapses and to differentiate between response and failure within the average trace of a selected synapse (Figure 4b, top). The given example in Fig. 4 illustrates the impact of noise in fast pulse trains. Even though a very small threshold multiplier of two is applied to the raw average trace the responses to the stimulus given at frames 130 and 140 are not detected. Moreover, the noise in the raw data (Figure 4b, top) obscures these responses, while the denoised trace (Figure 4b, bottom) reveals clear and distinct responses upon each of the five stimuli. Thus, the failure rate can be determined to be 0% after applying denoising with Neuroimage Denoiser (threshold multiplier 2), contrary to 40% in the raw trace (threshold multiplier 2). Additionally, when computed with a multiplier of two (yellow dashed line, Figure 4b), the low threshold leads to the detection of false positive responses in the raw trace. Quantitative analysis of the recordings demonstrated significant improvements in signal quality after denoising. The single-pixel SNR distributions (Figure 4c) show a marked shift toward higher values in the denoised data compared to the raw recordings. Peak analysis revealed a 2.00 ± 0.12-fold improvement in SNR (mean ± SEM, p < 0.001, paired t-test). Importantly, the denoising process preserved the original signal amplitudes, as evidenced by the strong correlation between raw and denoised measurements (R = 0.9998, p < 0.0001; Figure 4d).

**Figure 4.**
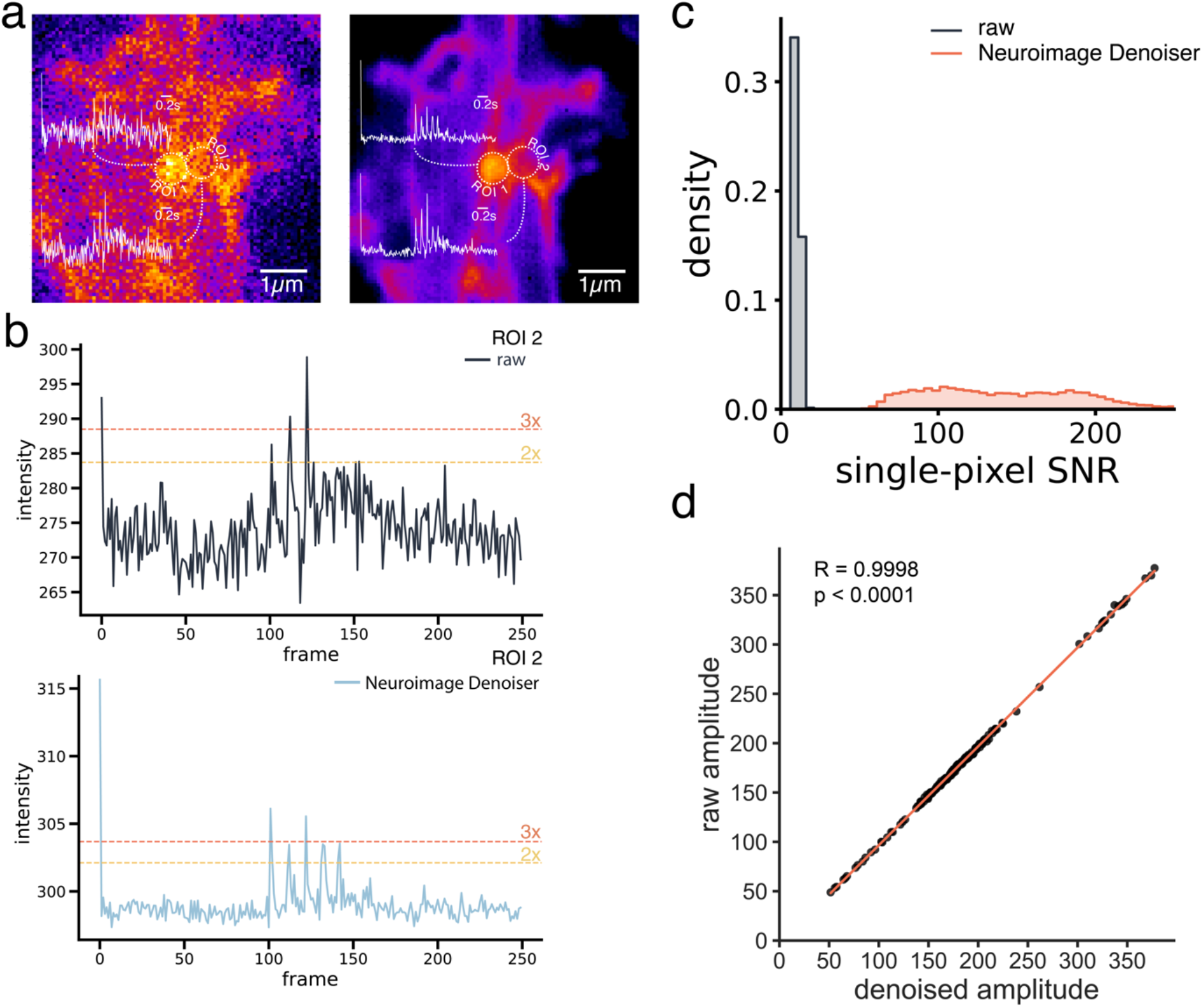
a) A crop of the recording with iGluSnFR.S72A^14^. The left panel represents the raw data with high noise levels, while the right panel shows the denoised data using Neuroimage Denoiser. b) Average intensity trace for ROI 2 over the recorded 250 frames. Stimulation is applied at frames 100, 110, 120, 130, and 140. The top graph is extracted from the raw recording and is characterized by significant noise and fluctuations. The bottom graph shows the denoised average intensity trace from ROI 2, where the noise level is substantially reduced, allowing clearer identification of responses to stimuli. Particularly, responses to the stimulation in frames 130 and 140, which are within the noise level in the raw trace, become visible in the denoised trace. The threshold for two low threshold multipliers is indicated as yellow and orange dashed lines for multipliers two and three, respectively. c) Single-pixel SNR distributions for raw and denoised recordings d) Correlation between raw and denoised signal amplitudes.

Next, we tested whether Neuroimage Denoiser can denoise recordings of pAAV-hSyn-synaptophysin-GCamp6f that emits fluorescence upon the binding of Calcium. Although this version of GCamp6f is also synaptically located, it shows far slower dynamics than the two iGluSnFR versions used in this study. Our results demonstrate that Neuroimage Denoiser effectively preserves GCaMP6f signals across different stimulation intensities, from 1AP to 5APs (Figure 5a). The denoised fluorescence traces closely match the temporal profile of the raw signals while substantially reducing noise, as evidenced by the smoother baseline and improved signal clarity (Figure 5a). Quantitative analysis revealed a significant improvement in single-pixel SNR in the denoised recordings (Figure 5b), showing an average 4.47 ± 0.23-fold improvement (mean ± SEM, p < 0.0001, paired one-sided t-test) for the synaptic responses.

**Figure 5.**
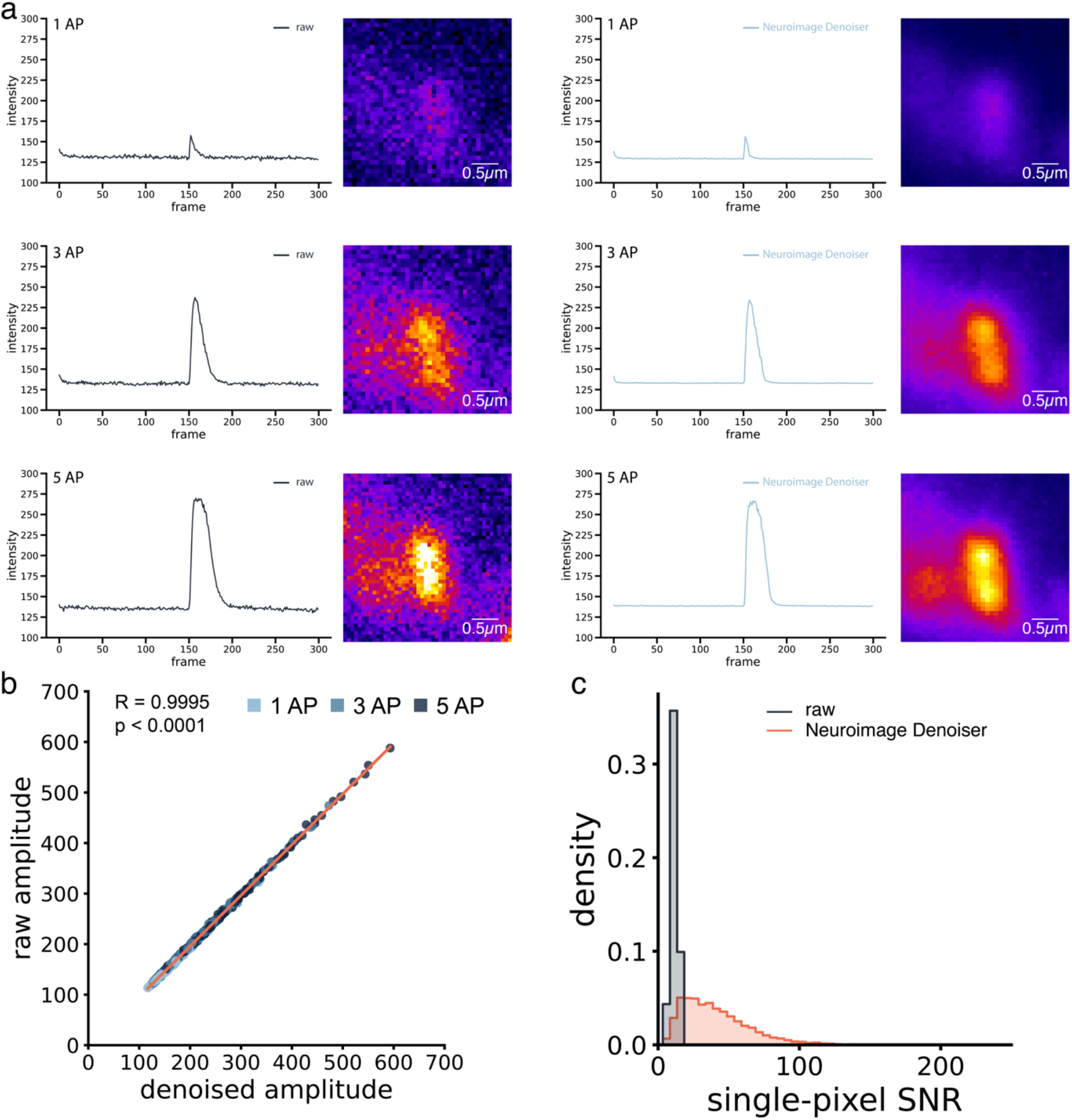
a) GCaMP6f responses to 1, 3, and 5 action potentials (APs) showing raw recordings (left) and corresponding denoised frames by Neuroimage Denoiser (right). Both fluorescence traces and peak response frames demonstrate signal preservation across different response magnitudes. b) Single-pixel SNR distribution for raw and denoised recordings. c) Correlation between raw and denoised signal amplitudes demonstrates high-fidelity preservation of response magnitudes (R = 0.9995, p < 0.0001)

Importantly, the denoising process maintained the original signal amplitudes, showing a very high correlation between raw and denoised measurements (R = 0.9995, p < 0.0001; Fig. 5c). This robust preservation of signal integrity was consistent across all stimulations (1, 3 and 5 Aps).

## Discussion

In this manuscript, we presented Neuroimage Denoiser – a U-Net-based^23^ deep-learning framework to reduce noise levels in microscopic imaging recordings. By incorporating the concepts of Noise2Noise^28^, the Neuroimage Denoiser framework is highly flexible for a wide array of experimental settings and does not require re-training for changed parameters. The Neuroimage Denoiser uses a self-supervised training approach without requiring noise-free ground truth. By automatically selecting training data on responsive structures like synapses or somata (on both spatial and temporal dimensions), the framework avoids overrepresenting background regions.

In the past, researchers developed several sensors to study functional neurobiology for the dynamics of calcium^30^, GABA^31^, dopamine^32^, acetylcholine^33^, and glutamate^13,31^ on a single synapse level. These recordings are superimposed with noise regardless of the sensor and acquisition rate. Consequently, a unified, flexible model capable of denoising recordings with the same localization would considerably ease the downstream processing.

Our comprehensive analysis demonstrates the framework’s effectiveness through both SNR improvements for peaks in responding synapses and single-pixel SNR distributions throughout the entire field of view. We tested our model, trained solely on iGluSnFR3.v857.SGZ^13^ data acquired at 100Hz, by applying the denoising routine to changed parameters, i.e., recording frequency (200, 500, and 1000 Hz; Figure 3) and sensor (iGluSnFR.S72A^14^ and GCaMP6f; Figure 4 and Figure 5), and obtained convincing results. Thus, the framework provides out-of-the-box utility for an array of changing experimental parameters and drastically reduces the need for re-training.

The proposed framework is capable of significantly reducing the noise in experiments with fast dynamics, as typical for glutamate imaging. One of the advantages of Neuroimage Denoiser is its ability to preserve the amplitude of synaptic responses, as evidenced by a high correlation between denoised and raw responses (Figure 2c, Figure 3f, Figure 4d, and Figure 5b). In all tested conditions, namely, changed recording frequency and the iGluSnFR3.v857.SGZ^13^ or GCamp6f sensor, the SNR for the responses was significantly improved by 5 to 9-fold depending on the observed noise level in the raw recordings. Only the iGluSnFR.S72A^14^ showed a lower average 2-fold improvement in SNR, likely because of the comparably weak response amplitudes. This consistent performance across different sensors and experimental conditions underscores the framework’s robustness and flexibility, which is a key strength of single-frame denoising approaches. Another important aspect is that this improved signal-to-noise level helps to resolve complex synapses spatially (Figure 3), can ease the analysis of spontaneous activity, and can be used as a functional readout for molecular manipulation to define release mechanisms in specific synapses.

We anticipate that Neuroimage Denoiser can be adopted in many laboratories as it has minimal hardware requirements, and the preparation of the training data is automatically balanced for optimal results. The framework is easy to use for data preparation, training, grid search, and inference (denoising) and does not require additional programming knowledge, making it accessible to a broad range of researchers in the field of neuroscience. Although glutamate imaging as a neural recording method can provide rich and diverse datasets, these data have a high complexity with many disruptive elements, such as noise, bright sensor depositions, or vesicular transport of sensor, that make the creation of ground truth data for training a model on responding regions of a recording challenging. We tested several potential training parameters (Table 1) to select the most performant parameter combination. However, users can freely choose a combination suited for their experimental set-up and analysis.

While initially developed for fast glutamate imaging responses, Neuroimage Denoiser has proven to be a versatile framework. Its successful application to both fast glutamate dynamics and slower calcium signals, along with consistent improvements in single-pixel SNR and SNR for responses across different experimental conditions, suggests it could serve as an out-of-the-box tool for enhancing the quality of various functional imaging experiments in neuroscience.

## Code availability

The code is available at https://github.com/s-weissbach/neuroimage_denoiser.

## Data availability

Data used for the training of Neuroimage Denoiser is available on Zenodo https://zenodo.org/records/14213082.

## Acknowledgment

The authors acknowledge financial support from the Emergent Algorithmic Intelligence Centre funded by the Carl-Zeiss Foundation. We thank Anita Heine, Nathalie Philipp, and Corinna Werkmann for technical support and excellent hippocampal culture preparation. The authors thank Stanislav Sys for critical feedback and discussion. We thank Arun Chhikara for the cloning of the GCaMP6f construct used in this study.

## Authors contribution

SW and MH conceived the idea and conceptualized the work. MB, CA, AK, and MH conducted experiments. SW conceptualized and implemented the Neuroimage Denoiser framework. SW and JM implemented the data preparation routine. SW created visualizations. SW, MB, and MH wrote the manuscript. SG revised the manuscript. MH and SG supervised the project and acquired funding.

## Ethics statement

All procedures involving animals were conducted in accordance with the Guidelines for the Care and Use of Laboratory Animals and approved by the District administration Mainz-Bingen (license number 41a/177-5865-§11 ZVTE, 30.04.2014). Animal handling complied with European Union guidelines (Directive 2010/63/EU) and local government (Rheinland-Pfalz, Germany) regulations for animal welfare (Tierschutzgesetz §4). No in vivo experiments were conducted.

