## Supplementary Figure for "Neuroimage Denoiser - a Deep Learning Framework for Removing Noise from Transient Fluorescent Signals in Functional Imaging"

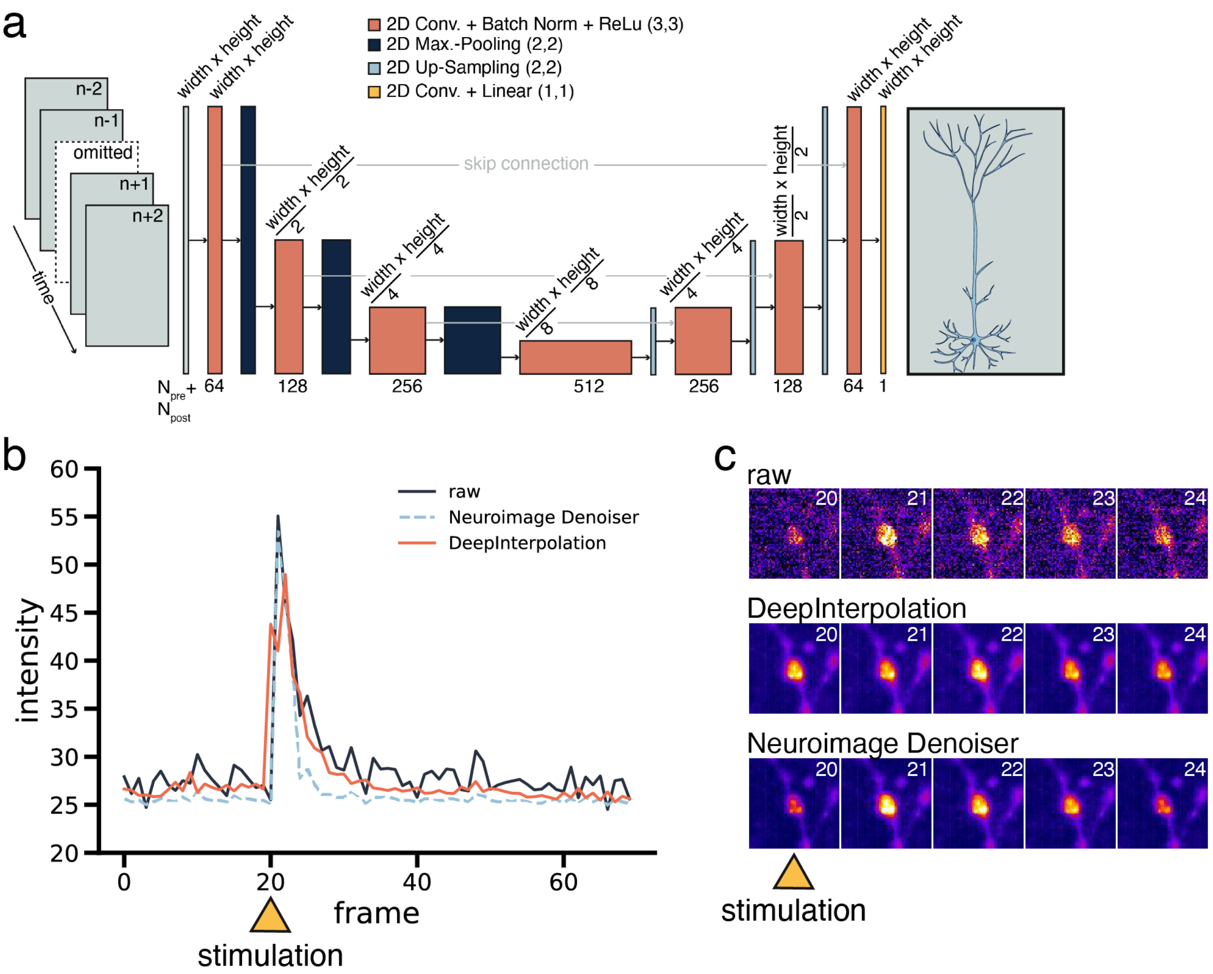


*Supplementary Figure 1 a) Schematic representation of a U-Net in the style of DeepInterpolation. Predictions are made based on five frames before and after the target frame (which is omitted). The model predicts tries to predict the omitted frame. b) Example of an artifact produced by a DeepInterpolation-styled approach (orange) to denoising for the fast kinetic of iGluSnFR. The peak is not preserved, instead, a double peak emerges with the original peak being of lower intensity than the preceding and succeeding frames. c) Raw, DeepInterpolation, and Neuroimage Denoiser denoised recordings of the synapse. Stimulation was applied at frame 20. It is visible that DeepInterpolation shifts the peak response from frame 21 to frame 22.*
